# ObiWan-Microbi: OMERO-based integrated workflow for annotating microbes in the cloud

**DOI:** 10.1101/2022.08.01.502297

**Authors:** Johannes Seiffarth, Tim Scherr, Bastian Wollenhaupt, Oliver Neumann, Hanno Scharr, Dietrich Kohlheyer, Ralf Mikut, Katharina Nöh

**Affiliations:** Institute of Bio- and Geosciences, IBG-1: Biotechnology, Forschungszentrum Jülich GmbH, Jülich, Germany; Computational Systems Biotechnology (AVT.CSB), RWTH Aachen University, Aachen, Germany; Institute for Automation and Applied Informatics, Karlsruhe Institute of Technology, Eggenstein-Leopoldshafen, Germany; Institute for Advanced Simulation, IAS-8, Forschungszentrum Jülich GmbH, Jülich, Germany; Microscale Bioengineering (AVT-MSB), RWTH Aachen University, Aachen, German

## Abstract

**Summary:** Reliable deep learning segmentation for microfluidic live-cell imaging requires comprehensive ground truth data. *ObiWan-Microbi* is a microservice platform combining the strength of state-of-the-art technologies into a unique integrated workflow for data management and efficient ground truth generation for instance segmentation, empowering collaborative semi-automated image annotation in the cloud.

**Availability and Implementation:** *ObiWan-Microbi* is open-source and available under the MIT license at https://github.com/hip-satomi/ObiWan-Microbi, along documentation and usage examples.

**Supplementary information:** Supplementary data are available online.

## Introduction

The combination of automated live-cell microscopy and microfluidic lab-on-chip technology unlocks unique spatio-temporal insights into the morphology of individual cells, the development of microbial populations, and community interactions (Burmeister and Grünberger, 2020). Time-lapse imaging produces large amounts of image data, calling for reliable high-throughput image processing pipelines. Deep learning approaches have become state-of-the-art for many bioimage applications (Moen *et al*., 2019), especially when adaptability to various morphologies and imaging modalities are required (Caicedo *et al*., 2019). Still, extracting microbial single-cell properties like cell count or area from microscopic image sequences remains challenging (Jeckel and Drescher, 2021).

The practical application of deep learning for segmentation (DLS) is often hampered by the lack of sufficient high-quality ground truth (GT) annotations for training and validation (Laine *et al*., 2021), as well as a limited accessibility for users due to complex software dependencies or special hardware requirements (Moen *et al*., 2019). Recently, solutions addressing individual aspects have been developed, such as Omnipose providing a user interface for cell segmentation (Cutler *et al*., 2021), or ImJoy that simplifies DLS usage utilizing browser-based user interfaces, a flexible plugin architecture and remote computation resources (Ouyang *et al*., 2019). Still, there is no integrated end-to-end workflow that provides state-of-the-art DLS methods for microbial live-cell images, is extensible, allows automated instance segmentation and convenient manual correction, connects to remote computational resources, is collaboration-ready, and integrates image and meta-data management.

To leverage integrated, efficient and reproducible workflows in microbial DLS with all the mentioned features optimized for fast GT creation, we developed *ObiWan-Microbi* (Fig. 1). *ObiWan-Microbi* comprises an extensible collection of DLS methods to be executed on a remote server, is customizable by end-users, comes with a user-friendly web app interface providing semi-automated DLS for image data stored in an OMERO backend (Allan *et al*., 2012), and facilitates team and project-based collaboration using cloud technology.

**Figure 1.**
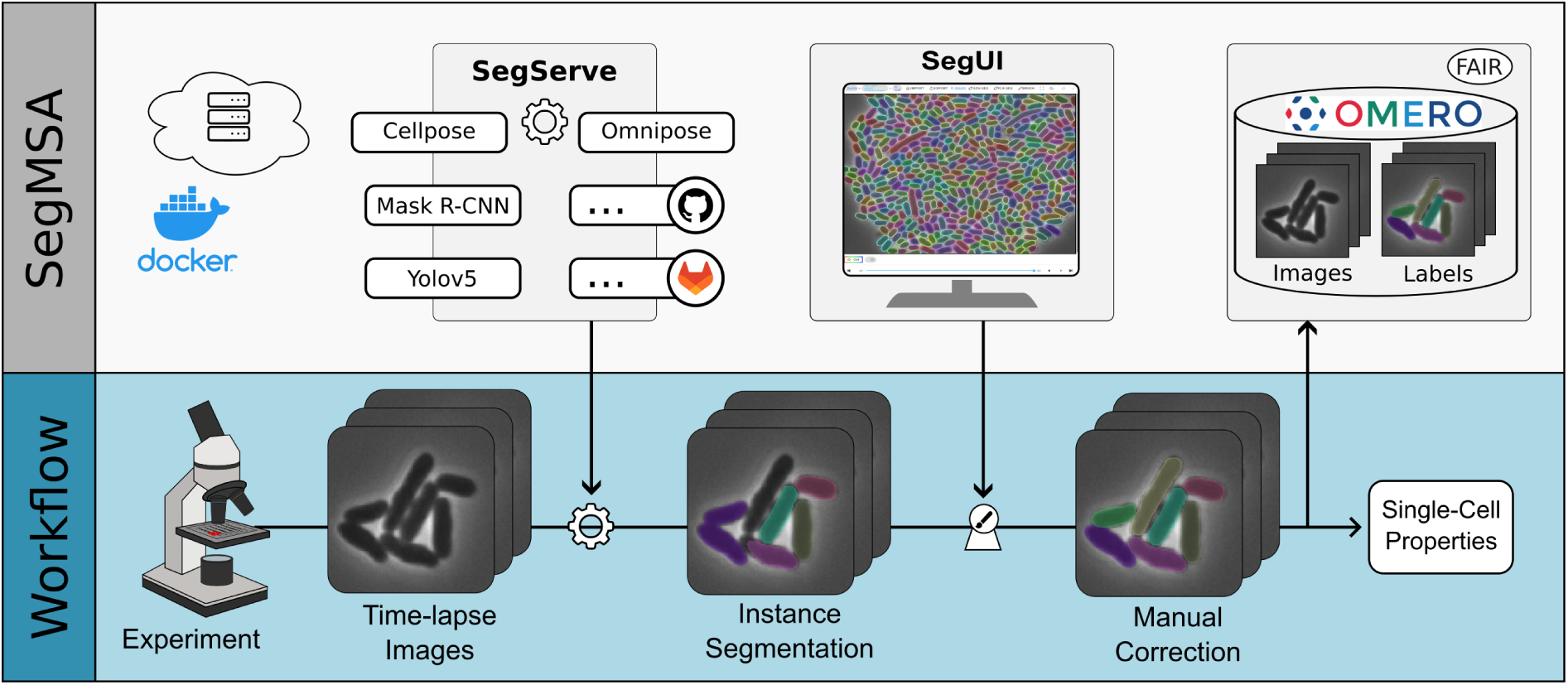
*ObiWan-Microbi* platform overview. Fully integrated end-to-end workflow for semi-automated image annotation and segmentation of time-lapse microscopy in the browser. *ObiWan-Microbi* consists of *SegServe, SegUI*, and OMERO jointly distributed in the microservice architecture *SegMSA*.

## Approach and Implementation

The open *ObiWan-Microbi* platform consists of three major parts (Fig. 1): *SegServe* for convenient DLS execution, the graphical user interface *SegUI*, and the microservice architecture *SegMSA* for deployment and integration of OMERO data management. DLS execution is often difficult in practice due to hardware and software requirements. *SegServe* is a convenient REST API designed to execute DLS methods, bundled in git repositories providing reproducible code execution (SI S.1). The execution is performed on a single server, accelerated by GPU hardware and batch processing, with no need for local installation or software dependency handling on the user-side. The REST API functionality is automatically documented using FastAPI.

To unlock the power of DLS for GT generation, we developed the user-friendly graphical user interface *SegUI*, an Angular and Ionic based web app that connects to the OMERO data backend. *SegUI* allows browsing image data and performing fully manual or fast semi-automated image annotation of time-lapse sequences. Results from automated DLS by *SegServe* are available in the web app *SegUI* for manual correction using efficient drawing tools, a data model to undo/redo annotation actions, and multi-label annotations for complex imaging data. Custom DLS approaches are supported by *SegUI* to readily include new or optimized segmentation approaches and parameters settings. Thus, *SegUI* provides a convenient and interactive DLS experience for collaborative microfluidic image annotation.

*SegUI* and *SegServe* are combined with OMERO to form the microservice architecture *SegMSA* (SI S.2). OMERO takes care of image data handling and segmentation storage including user and access management, facilitating FAIR data principles. This way, OMERO-compatible modules, such as Fiji can be utilized to extract single-cell properties. *SegMSA* utilizes docker to manage services and their connections providing a convenient installation procedure for the microservice architecture deployment.

## Results

The combination of the three components forms the platform *ObiWan-Microbi*, used to segment microbial organisms including *E. coli, C. glutamicum* and *B. subtilis* (SI S.3). For semi-automated segmentation, four pre-trained DLS algorithms are provided out-of-the-box and accessible in *SegUI* including Cellpose (Stringer *et al*., 2021), Omnipose (Cutler *et al*., 2021), Mask R-CNN trained on simulated image data (Sachs *et al*., 2022) and Yolov5 for general object detection in images. Using semi-automated segmentation, an annotation speed of more than 200 cells/minute was achieved (SI S.4).

## Conclusion

*ObiWan-Microbi* is a collaborative annotation platform for the efficient creation of high-precision GT data that debottlenecks the broad usage of DLS in microbial image processing, especially for distributed teams. Based on growing GT data sets, and in combination with integrated training workflows (Scherr *et al*., 2022), we are looking forward to seeing increased usage of DLS methods and publicly available benchmarks for training and validation in the domain of live-cell imaging.

## Acknowledgement

JS and KN acknowledge the inspiring scientific environment provided by the Helmholtz School for Data Science in Life, Earth and Energy (HDS-LEE) and thank Wolfgang Wiechert for continuous support.

