## Supplementary Information for "ObiWan-Microbi: OMERO-based integrated workflow for annotating microbes in the cloud"

### S.1 Expanding DLS model zoo: Executor repository guidelines

State-of-the-art DLS methods develop and improve rapidly. Therefore, *ObiWan-Microbi* allows using an extensible zoo of DLS methods, enabling highly customized models or completely new approaches to be used in the platform. The extensibility is achieved by packaging DLS methods into public git repositories that are accessed via a unique URL. The URL and specific method parameters are sufficient for *SegUI* and *SegServe* to perform DLS on image data using remote computation resources while maintaining reproducibility. To allow DLS execution remotely through *SegServe*, the repository must contain code to perform DLS inference and has to follow a special structure to define software dependencies and entry points. Examples for such a repository layout are available in our three exemplary DLS method repositories[1,2,3] and a detailed description about adding your own custom model executor is provided in the *SegServe* repository[4].

[1] https://github.com/hip-satomi/MMDetection-Executor
[2] https://github.com/hip-satomi/Cellpose-Executor
[3] https://github.com/hip-satomi/Yolov5-Executor
[4] https://github.com/hip-satomi/SegServe

### S.2 Microservice architecture

Microservice architectures make use of network interfaces between various services while providing good opportunities for scaling to large numbers of users, for example, using load balancers. Well-defined interfaces allow using various different programming languages and configurations across the services and, therefore, taking optimal choices for specific use cases is possible.

By its modular design, it is straightforward to add custom microservices that integrate missing functionality into the existing system. Thus, *ObiWan-Microbi* benefits from the existing OMERO and OMERO Web services and adds *SegUI* as well as *SegServe* to build the microservice architecture *SegMSA* (Figure S.1).

The front end connections are handled by a reverse nginx proxy that redirects requests based on their URL to the target services. Therefore, existing OMERO functionality can be used out-of-the box. This functionality is augmented with the additional features of *ObiWan-Microbi*. In combination with the docker containerization, installation and administration of the *ObiWan-Microbi* microservice architecture is straightforward, for example, the initial installation can be performed with only three command lines on a Linux computer.

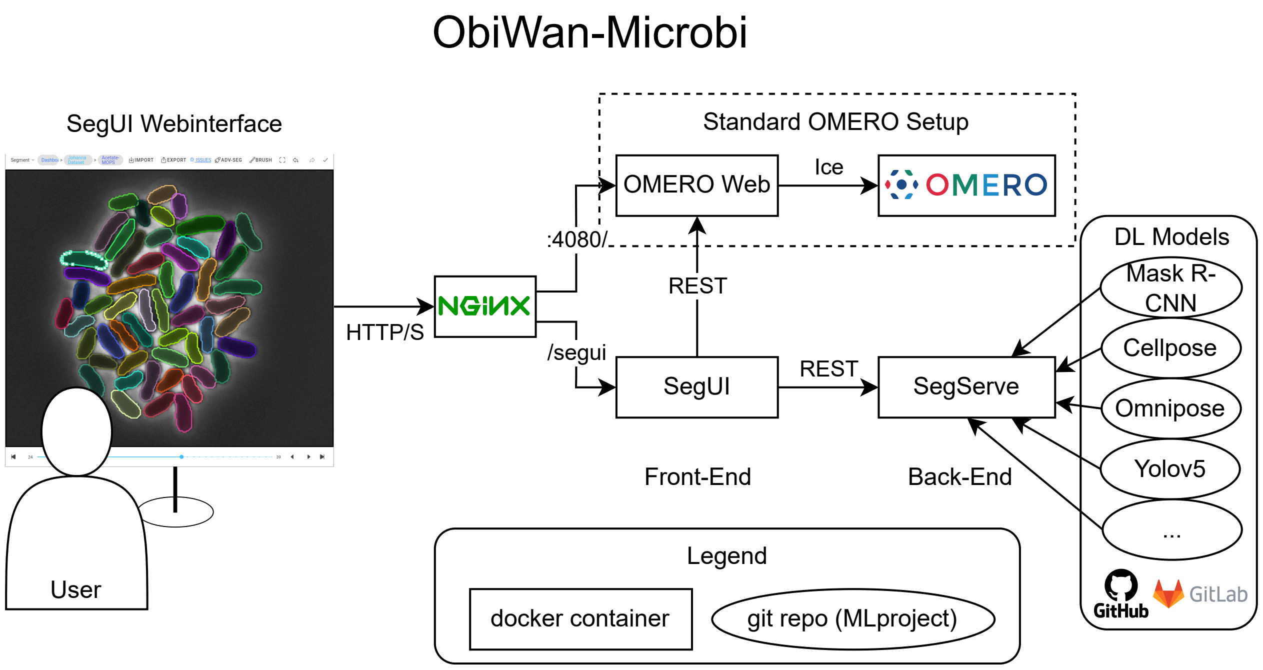


Fig. S.1: Microservice architecture design of *ObiWan-Microbi*. Docker containers are connected via network connections communicating using network protocols such as REST, HTTP/S and Ice. Access to OMERO is managed using the REST API provided by OMERO Web. *SegServe* installs deep learning methods on-demand from git repositories.

### S.3 Application of deep learning methods to microfluidic live-cell image data: *E. coli*, *C. glutamicum* and *B. subtilis*

For qualitative testing of *ObiWan-Microbi*'s deep learning segmentation (DLS) methods, we randomly selected phase contrast images from microfluidic experiments featuring three different microbial organisms, namely *Escherichia coli*, *Corynebacterium glutamicum*, and *Bacillus subtilis* grown in monolayers. The microfluidic single-cell cultivation including the live-cell imaging setup are similar to the one described by Kaganovitch (2018). Whereas *E. coli* and *C. glutamicum* show a classic rod-shape morphology, *B. subtilis* grows into filamentous-like structures. In Figure S.2 selected phase contrast images are shown along with the corresponding segmentation results produced by three state-of-the-art DLS methods: *Cellpose* (Stringer, 2021) and *Omnipose* (Cutler, 2021) are used with their default parameters and pre-trained weights that came with their implementation. *Mask R-CNN* is trained on simulated data using adapted hyperparameters (Sachs, 2022). All model executors are publicly available in our github repositories[5,6].

From Figure S.2, we observe that *Omnipose* performs best and is the only method segmenting *B. subtilis* microbes properly. This can be explained by the design of the method addressing segmentation of filamentous morphologies and the large size of the training data set including 46,000 manually annotated microbial cell instances. *Cellpose* only detects very few cells across all three morphologies with consistently too large instance masks. One reason for the deficient segmentation quality might be the training data consisting of biomedical images with dense cell colonies looking very different from the selected microbial images. The Mask R-CNN method trained on simulated data shows medium performance for *E. coli* and *C. glutamicum*, but fails to segment *B. subtilis*, likely because of a lack of filamentous morphologies in the training data and the difficulty for bounding box based detectors to deal with filamentous structures.

As a conclusion, we recognize that existing methods, in particular *Omnipose*, provide good baselines to utilize semi-automated annotation for ground truth creation and decrease the effort for manual annotation. Furthermore, not unsurprising, large annotated data sets for training comprising various morphologies lead to superior instance segmentation performance. Using *ObiWan-Microbi* it is now possible to integrate custom pre-trained DLS methods (see section S.1) and conveniently create these large data sets.

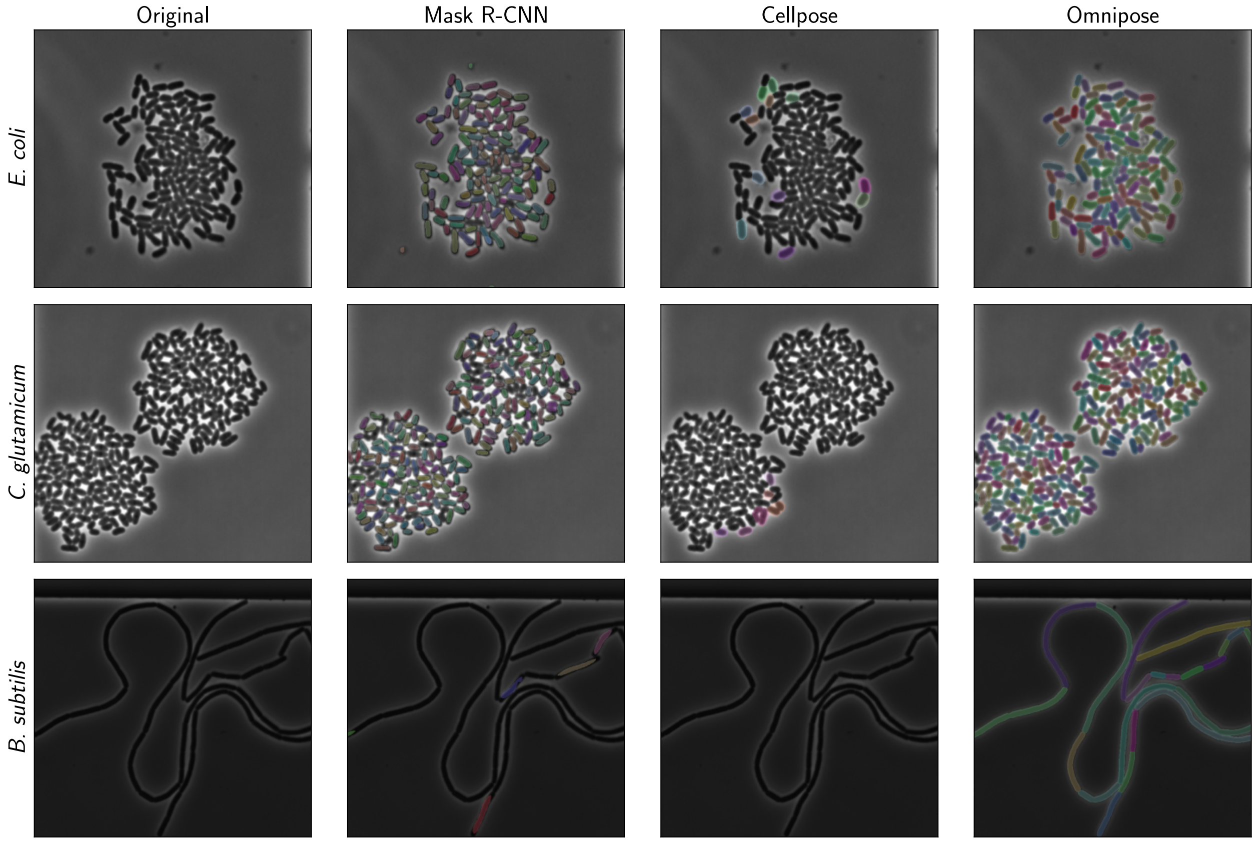


Fig. S.2: Segmentation results of microfluidic phase contrast images showing *E. coli*, *C. glutamicum* and *B. subtilis* bacteria using *Mask R-CNN*, *Cellpose* and *Omnipose* DLS approaches. Images are screenshots from *SegUI* in the browser. Segmented cell instances are visualized with bright colors.

[5] https://github.com/hip-satomi/MMDetection-Executor

[6] https://github.com/hip-satomi/Cellpose-Executor

### S.4 Semi-automated annotation speed

The semi-automated annotation of *ObiWan-Microbi* is designed to annotate high-quality ground truth data at scale utilizing pre-trained DLS approaches. To demonstrate these capabilities we selected four phase contrast frames of a growing *C. glutamicum* colony (Schito, 2022) and performed the segmentation using Omnipose (Cutler, 2021). Artifacts (often due to the chamber structures), under-segmentations (multiple cells merged into one annotation) and missing cell detections are manually corrected in the *SegUI* user interface (see Figure S.3). A single annotator performed this process for the four frames containing 2,478 cell instances in 10 minutes (604 s) resulting to an annotation speed of 246 cells/min.

The annotation speed is mainly determined by the already good performance of the Omnipose DLS approach. For other microbial morphologies and imaging setups the annotation speed can vary and a more detailed comparison of different annotation tools on a broad range of biological image data is needed in the future. Nonetheless, the example shows that well-trained DLS approaches are suited for the human-in-the-loop creation of GT data in *SegUI*. Thus, the *ObiWan-Microbi* platform is a step towards large and multi-morphological data sets containing thousands of annotated cell instances.

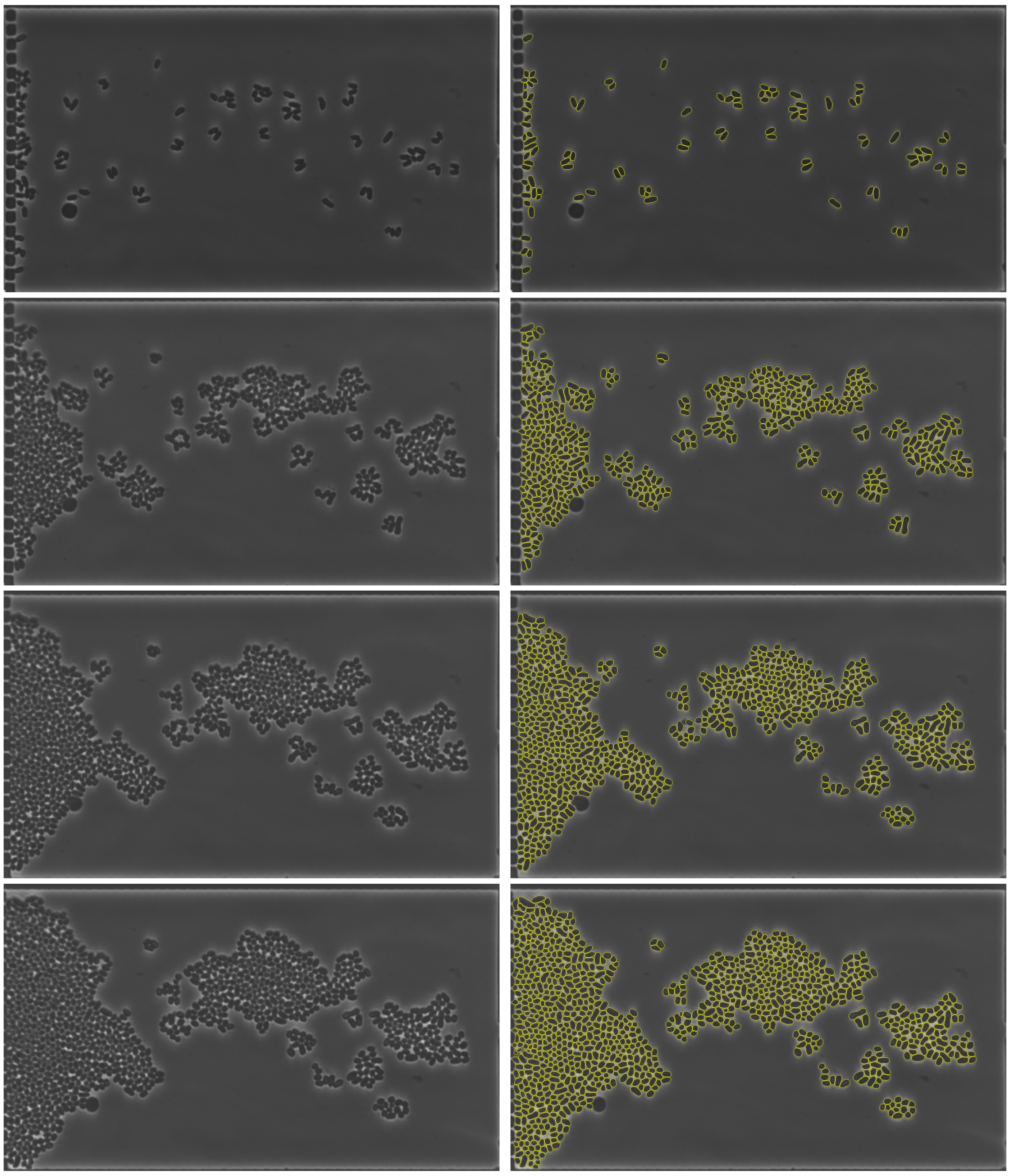

Fig. S.3: Microscopic phase contrast images (left) and overlayed yellow contours of instance annotations (right) for 4 time points of a growing *C. glutamicum* colony. For cultivation details see Schito (2022).
